# PGC-1α and PPARs cooperatively mediate photoreceptor neuroprotection in *rd1* mouse inherited retinal degeneration

**DOI:** 10.1101/2025.10.21.683632

**Authors:** Lan Wang, Jie Yan, Javier Sancho-Pelluz, Qianlu Yang, Zhulin Hu, Kangwei Jiao, Mathias W. Seeliger, François Paquet-Durand

## Abstract

Retinitis pigmentosa (RP) is a group of inherited diseases characterized by a primary rod photoreceptor dysfunction and progressive rod and cone cell death. Due to their very high energy demand, the degeneration of photoreceptors may be linked to insufficient energy supply or metabolic imbalance. Critical transcription factors that regulate metabolism such as peroxisome proliferator-activated receptors (PPARs) and their co-activator PGC-1α have been found to play important roles in neurodegenerative diseases, but their potential roles in RP have yet not been disclosed. In this study, we used organotypic retinal explant cultures derived from the *rd1* mouse model for RP to investigate the effects of PPARα, PPARγ, PPARβ/δ agonists, as well as PGC-1α activation and inhibition. Photoreceptor death in the outer nuclear layer (ONL) of the retina was quantified using the TUNEL assay, while *in situ* activity assays were used to monitor effects of PPARs and PGC-1α on poly (ADP-ribose) polymerase (PARP) and calpain activity. In addition, we performed immunostainings to evaluate poly (ADP-ribose) (PAR) generation and activation of calpain-1 and calpain-2. We found that PPARβ/δ agonists had limited effects, while activation of PPARα, PPARγ, and PGC-1α significantly reduced photoreceptor death and PARP activity in *rd1* retina. Conversely, inhibition of PGC-1α had a strong detrimental effect on photoreceptor viability. Activation of the histone deacetylase sirtuin-1, an upstream agonist of PGC-1α, had no effect unless it was combined with simultaneous inhibition of PARP. Furthermore, PPARγ and PGC-1α effectively suppressed overall calpain activity and overactivation of calpain-2, alleviating photoreceptor degeneration caused by Ca^2+^ imbalance. In summary, our data supports the concept of a PARP–sirtuin-1–PGC-1α–PPAR–PARP feedback control that connects defective energy metabolism to photoreceptor degeneration. Specifically, our findings suggest that PPARα, PPARγ, and PGC-1α cooperate to preserve photoreceptor viability, highlighting PPAR-signaling as a promising target for future therapeutic interventions.

## Introduction

Retinitis pigmentosa (RP) is an inherited retinal degeneration (IRD) that affects approximately 1:4.000 people, with more than 2 million patients worldwide^1^. Initially, causative genetic mutations lead to the degeneration of rod photoreceptors and night blindness. As the disease progresses, a secondary degeneration of cone photoreceptors sets in, resulting in progressive visual field constriction – so-called tunnel vision – and ultimately loss of central vision and complete blindness^2^.

The mechanisms of RP photoreceptor degeneration are only incompletely understood. While apoptosis was initially thought to be the primary pathway^3^, emerging evidence highlights the role of alternative cell death mechanisms. For instance, photoreceptor cell death has been suggested to be caused by cyclic guanosine monophosphate (cGMP)-mediated Ca^2+^ overload and over-activation of Ca^2+^-dependent calpain-type proteases^4, 5^. Experimental evidence also points to an involvement of poly (ADP-ribose) polymerase (PARP) and excessive formation of poly (ADP-ribose) (PAR) polymers. Since PARP consumes high levels of nicotinamide dinucleotide (NAD^+^), its overactivation has been linked to dysfunction of other NAD^+^-dependent enzymes, notably of sirtuin-type histone deacetylases critical for mitochondrial homeostasis ^6^.

Recent studies have highlighted links between RP and energy metabolism, with lipid and fatty acid metabolism likely playing an important role in its pathogenesis^7^. The renewal and regeneration of membrane discs in photoreceptor outer segments rely on continuous lipid synthesis, transport, and degradation^8^. Lipids participate in the regulation of phototransduction and visual cycle^9^. Given the specialized structure and function of photoreceptors, retinal energy requirements are extremely high, and fatty acid oxidation likely is important to satisfy these requirements^10^. Additionally, lipids and their metabolites may function as signaling molecules, influencing various cellular processes^11^.

Key players in lipid and energy metabolism are peroxisome proliferator-activated factors (PPARs), ligand-activated transcription factors belonging to the nuclear receptor family, including PPARα, PPARγ, and PPARβ/δ^12^. Long-chain fatty acids and arachidonoids serve as PPAR ligands, triggering the transcription of target genes upon PPAR binding^13^. PPARα regulates fatty acid oxidation in peroxisomes and mitochondria, helping to maintain energy homeostasis^14^. PPARγ is mainly associated with lipogenesis to reduce lipid levels^15^, and PPARβ/δ is implicated with fatty acid regulation and lipolysis^16^. Within the eye, PPARs have been related to pathologic processes in various retinal diseases, such as age-related macular degeneration (AMD) and diabetic retinopathy (DR)^17–21^.

Transcriptional activity and gene expression levels of PPARs are influenced by interactions with co-regulators such as PPARγ coactivator-1α (PGC-1α). PGC-1α is encoded by the *PPARGC1A* gene and forms complexes with PPARs to recruit additional cofactors, significantly enhancing their transcriptional activity on downstream target genes^22^. A key upstream regulator of PGC-1α is sirtuin-1, which can enhance PGC-1α activity through deacetylation, but its function may be compromised when excessive PARP activation depletes NAD^+^.

To investigate the roles of PPARs and PGC-1α in RP, we analyzed gene expression profiles and utilized organotypic retinal explant cultures derived from the retinal degeneration 1 (*rd1*) mouse, a well-established model of RP. Explants were treated with either subtype-specific PPAR agonists or a PGC-1α activator or inhibitor. We found that activation of both PPAR or PGC-1α can prevent *rd1* photoreceptor degeneration. This protective PPAR / PGC-1α signaling appears to be dependent on the activity of sirtuin-type deacetylases and can be counteracted by excessive PARP activity. Our data stresses the importance of PPAR / PGC-1α signaling for photoreceptor survival and highlights the corresponding up- and down-stream pathways as targets for therapeutic intervention.

## Results

### Expression patterns of PPAR and PGC-1α in the retina

An initial study of the GSE62020 *rd1* transcriptome dataset^23^, using MCODE cluster analysis^24^ and cytoHubba^25^ with the topological algorithms Degree, Closeness, Maximum Neighborhood Component (MNC), Edge Percolated Component (EPC) identified *Ppara* as a shared hub gene (Figure 1A). Remarkably, another hub gene found in this process was early growth response protein-1 (EGR1), which had already been identified in a recent single-cell RNA sequencing study performed on *rd1* and wild-type (*wt*) retina^26^. This provided for an independent validation of the hub gene identification and motivated us to further investigate the role of PPARα in the progression of *rd1* retinal cell death. We then analyzed the same dataset comparing *wt* and *rd1* retina at various post-natal (P) days. This transcriptomic data was related to the percentage of dying cells in ONL, as evidenced by the TUNEL assay, with *wt* data set as baseline for the comparative analysis. In *rd1* retinas, retinal cell death increased significantly from P9 to P13, and was still significantly elevated at P21 (Figure 1B). The expression of the *Ppara* gene encoding PPARα was markedly reduced at P11, *i.e.* at the onset of photoreceptor degeneration. Further analysis of other PPAR family members found a significant downregulation of *Ppard* (PPARβ/δ) expression only at P15, *i.e.* after the peak of ONL cell death. In contrast, *Pparg* (PPARγ) expression remained stable during the period of major photoreceptor loss up until P21 (Figure 1B).

**Figure 1.**
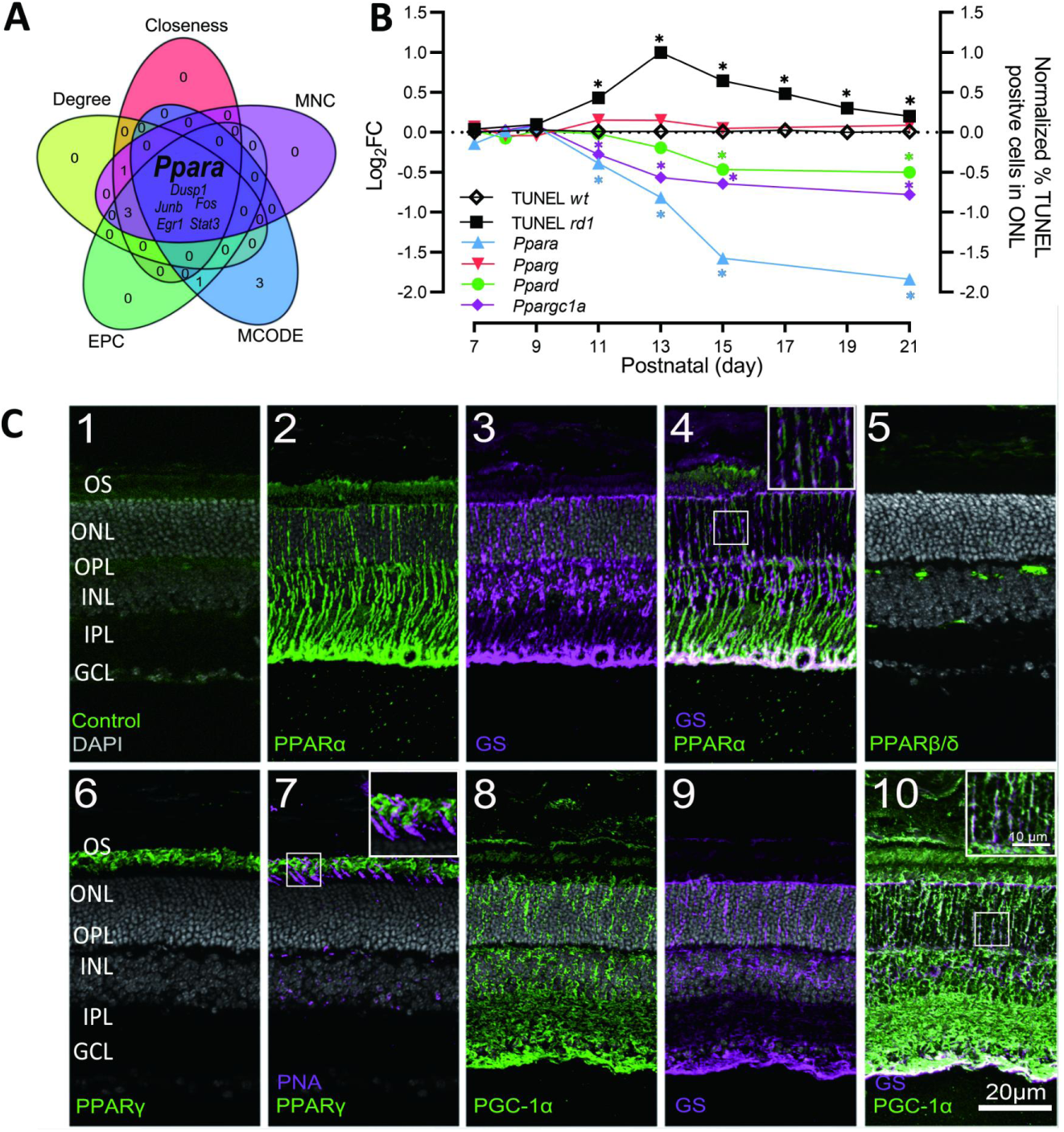
Analysis of PPAR and PGC-1α gene expression during *rd1* photoreceptor degeneration. **A**) cytoHubba analysis of *rd1* transcriptomes using Degree, Closeness, Maximum Neighborhood Component (MNC), Edge Percolated Component (EPC), and MCODE algorithms identified *Ppara* as a shared hub gene. **B**) Relative expression of *Ppara*, *g*, *d* and *Ppargc1a* genes (Log2FC; left y-axis) during the first 3 postnatal weeks. Retinal cell death expressed as normalized percentage of TUNEL positive cells in the outer nuclear layer (ONL) is shown for comparison (right y-axis). **C**) Immunostaining for PPARα, γ, β/δ and PGC-1α (green) at post-natal day (P) 30 in wild-type (*wt*) retina. DAPI (grey) was used as nuclear counterstain; negative control for secondary antibody shown in 1. Glutamine synthetase (GS; magenta) was used as Müller cell marker in 3 and 9; peanut agglutinin (PNA; magenta) was used to label cones in 7. Lack of colocalization between GS and PPARα or PGC-1α indicates that these are not expressed in Müller cells. Co-staining for PNA and PPARγ shows that PPARγ is expressed in rod outer segments (OS), but not in cones. Significance levels: * = *p* < 0.05. OPL = outer plexiform layer, INL = inner nuclear layer, IPL = inner plexiform layer, GCL = ganglion cell layer; scale bar = 20 µm.

Since mRNA expression does not necessarily represent protein levels, we used immunofluorescence on mature P30 *wt* retina to determine PPAR protein expression and localization. PPARα was strongly expressed across the retina, spanning from the photoreceptor outer segments (OS) all the way through to the ganglion cell layer (GCL) (Figure 1C2). To assess whether PPARα was expressed in Müller cells, we performed co-immunostaining with the Müller cell marker glutamine synthetase (GS). In the outer nuclear layer (ONL) PPARα signal did not colocalize with GS, suggesting that PPARα was not localized to Müller cells within the outer retina (Figure 1C4). Nevertheless, PPARα appeared to be expressed in Müller cell end feet within the GCL. PPARβ/δ signals were detected at the level of the outer plexiform layer (OPL) and at the inner border of the inner nuclear layer (INL), indicating an expression in inner retinal blood vessels (Figure 1C5). PPARγ was mainly expressed in the OS of photoreceptors (Figure 1C6). Co-staining of PPARγ with the cone marker peanut agglutinin (PNA) showed the PPARγ signal was localized outside of the PNA-positive regions, indicating predominant expression in rod photoreceptors (Figure 1C7).

To exert their functions, PPARs require transcriptional co-regulators such as PGC-1α, which strongly enhances PPAR-mediated transcriptional activity on target genes^22^. We thus investigated retinal PGC-1α gene expression and protein distribution patterns. Expression of *Ppargc1a* mRNA continuously declined concomitant with *rd1* photoreceptor loss (Figure 1A), suggesting that its deficiency may contribute to cell death. Immunostaining showed strong PGC-1α expression in the INL, inner plexiform layer (IPL), GCL, and photoreceptor layer (Figure 1C8). PGC-1α immunofluorescence did not colocalize with the Müller call marker GS (Figure 1C10).

### Effect of PPARs and PGC-1α on *rd1* photoreceptor death

To explore potential links between PPAR activity and photoreceptor degeneration, we treated *rd1* and *wt* organotypic retinal explants with the PPARα agonist GW590735 (abbreviated as GW735), the PPARγ agonist Inolitazone dihydrochloride (hereinafter called Inolitazone), and the PPARβ/δ agonist GW501516 (abbreviated as GW516). The effects of these treatments were assessed using the TUNEL assay to quantify cell death in the ONL, and dose-response curves were established for each agonist. In *wt* retinal explants, the number of TUNEL-positive cells in the ONL was low (Figure S1A, B; Table S1B). In untreated *rd1* explants, ONL cell death was significantly increased (Figure 2A, B; Table S1A). Treatment with GW735 and Inolitazone markedly reduced photoreceptor cell death in *rd1* retina and effective concentrations were well tolerated in *wt* retinas (Figure 2A, B; Figure S1A, B; Table S1A, B). However, treatment with GW516 did not lead to a significant reduction of TUNEL-positive cells in the *rd1* ONL, yet, at the same concentration, it significantly increased *wt* retina ONL cell death (Figure 2A, B; Figure S1A, B; Table S1A, B). Taken together, activation of PPARα and PPARγ appeared to significantly delay *rd1* photoreceptor degeneration.

**Figure 2.**
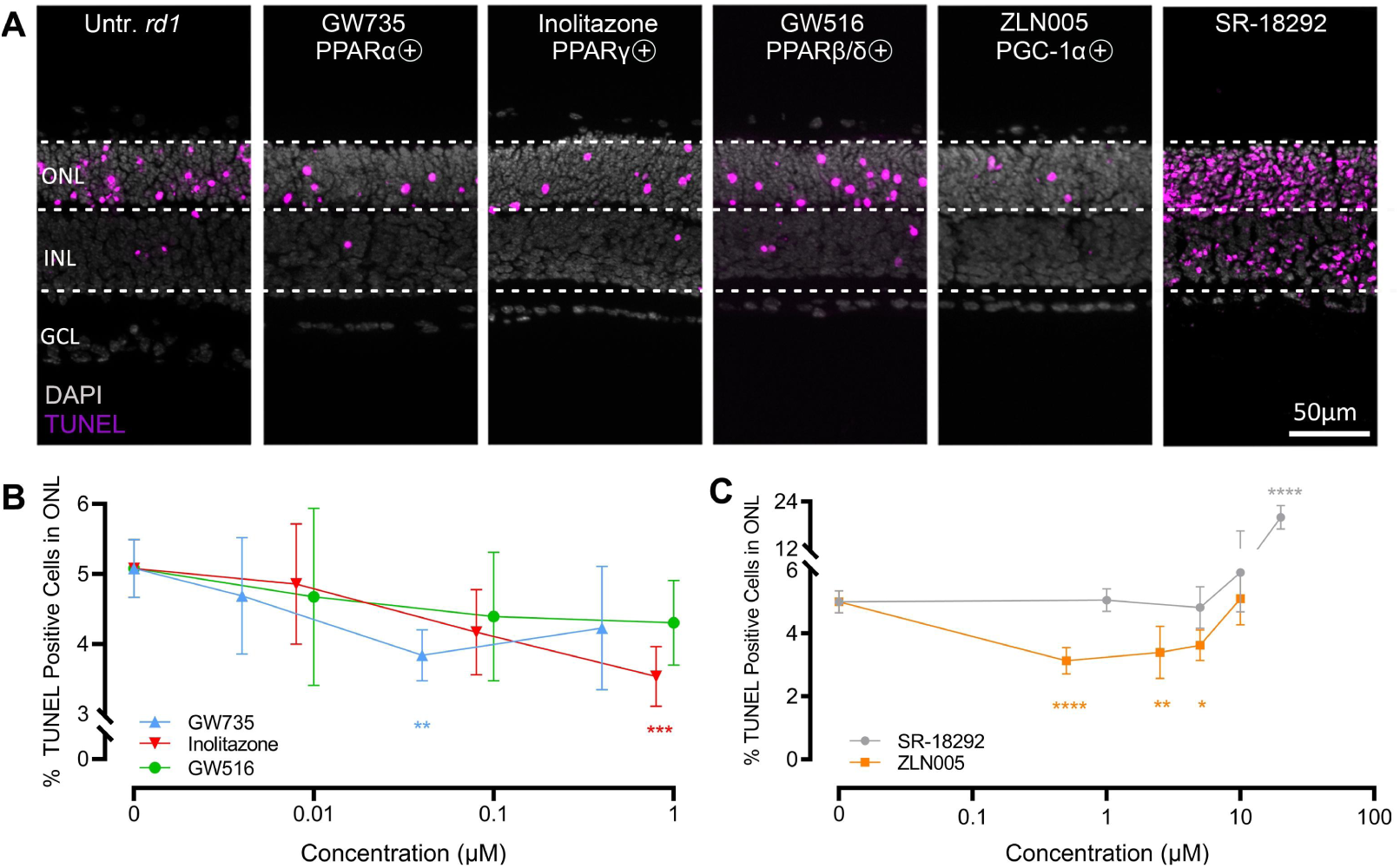
Photoreceptor cell death is decreased by PPAR and PGC-1α agonists. **A**) TUNEL assay labeling dying cells (magenta) in *rd1* mouse retinal explant cultures. DAPI (grey) was used as nuclear counterstain. Untreated (Untr.) *rd1* retina was compared to treatment with either 0.04 µM PPARα agonist GW735, 0.8 µM Inolitazone (PPARγ agonist), 1 µM GW516 (PPARβ/δ agonist), 0.5 µM ZLN005 (PGC-1α activator), or 20 µM SR-18292 (PGC-1α inhibitor). **B**) Dose-response curves for GW735, Inolitazone, and GW516 in *rd1* retinal cultures. In the outer nuclear layer (ONL), 0.04 µM GW735 and 0.8 µM Inolitazone, respectively, significantly reduced cell death. **C**) Dose-response curves for ZLN005 and SR-18292. ZLN005, at 0.5 to 5 µM, significantly reduced ONL cell death while 20 µM SR-18292 increased ONL cell death. Statistical testing: One-way ANOVA and Dunnett’s multiple comparisons test. Error bars represent SD; significance levels: * = *p* < 0.05; ** = *p* < 0.01; *** = *p* < 0.001; **** = *p* < 0.0001; INL = inner nuclear layer, GCL = ganglion cell layer; scale bar = 50 µm.

To functionally characterize the role of the PPAR coactivator PGC-1α, we applied the PGC-1α activator ZLN005 and the PGC-1α inhibitor SR-18292, respectively, to retinal *rd1* and *wt* retinal explant cultures. In *rd1* explants, SR-18292 significantly increased the number of TUNEL-positive cells in the ONL to 20.03 % (± 3.00 SD, n=3) (Figure 2A, C; Table S1A), whereas in *wt* retinas, it caused only 2.35 % (± 0.60 SD, n=5) cell death. (Figure S1A, B; Table S1B). This indicates that the effect of SR-18292 was not due to general cytotoxicity but instead was mediated through a specific mechanism in *rd1* retinas. Conversely, ZLN005 significantly reduced photoreceptor cell death in *rd1* retinas already at relatively low concentrations (Figure 2A, C; Table S1A).

### PGC-1α activates PPARα and PPARγ

The RNA-seq data indicated a significant decline in *Ppargc1a* and *Ppara* expression, suggesting a potential synergistic interaction (*cf*. Figure 1B). PGC-1α and PPARα were both expressed in the ONL, while PPARγ was detected in the OS, further supporting the idea that they might be functionally connected in photoreceptors (*cf*. Figure 1C). To study the role of PGC-1α in PPAR-mediated photoreceptor neuroprotection, we visualized the expression changes of PPARα and PPARγ in *rd1* retinal explants after treatment with the PGC-1α activator ZLN005 (Figure 3).

**Figure 3.**
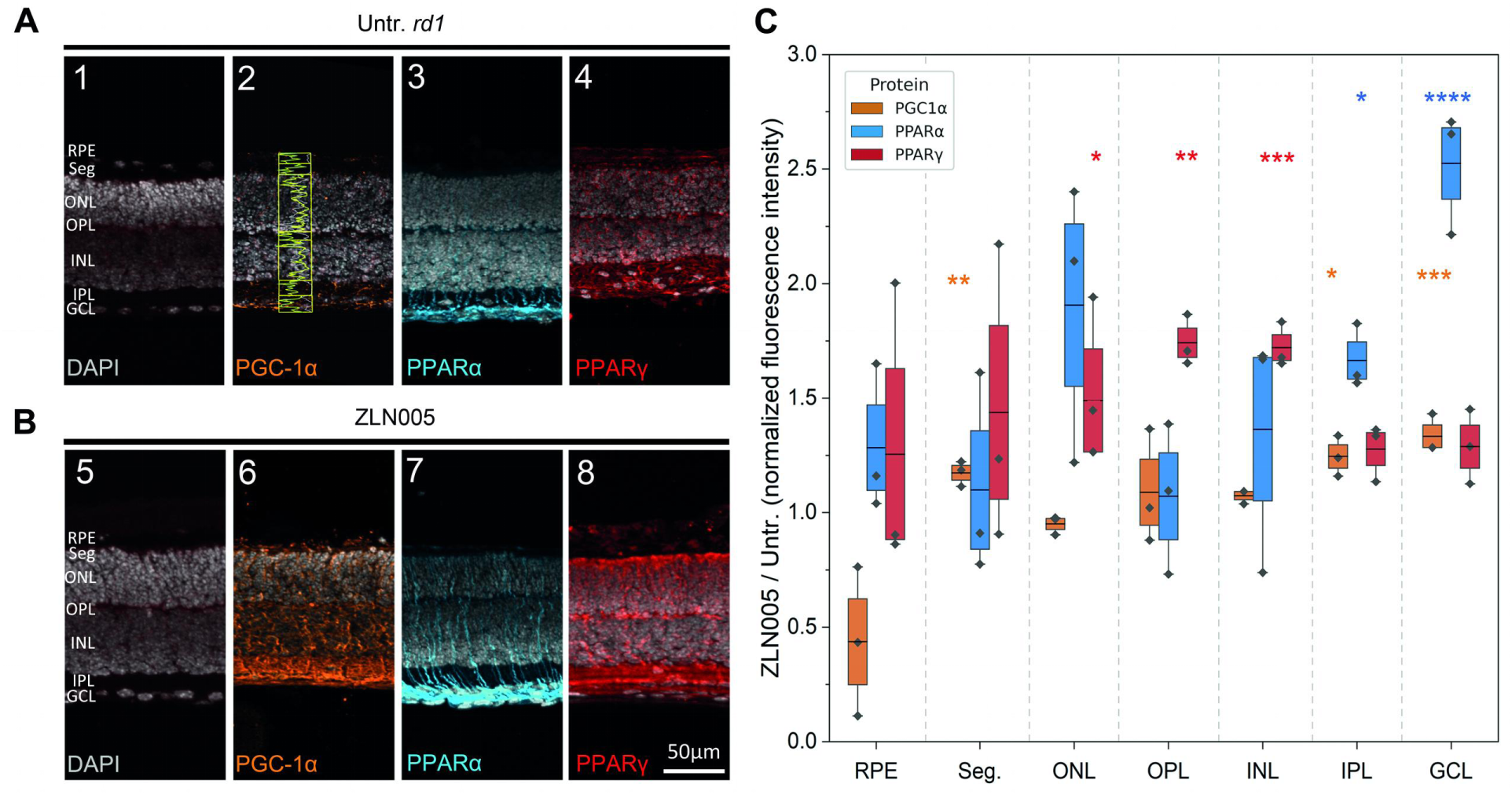
ZLN005 increases the expression of PGC-1α, PPARα, and PPARγ. **A**) Immunostaining for PGC-1α (orange), PPARα (cyan), and PPARγ (red) in untreated (Untr.) *rd1* retina at post-natal day (P) 11. DAPI (grey) was used as nuclear counterstain. **B**) Immunostaining for PGC-1α, PPARα, and PPARγ in P11 *rd1* retina treated with PGC-1α activator ZLN005. **C**) Box and whisker plot for fluorescence intensity levels of PGC-1α, PPARα, and PPARγ in retina treated with the PGC-1α activator ZLN005. Data normalized relative to untreated control for each retinal layer. Statistical testing: Student’s *t*-test. Error bars represent SD; significance levels: * = *p* < 0.05; ** = *p* < 0.01; *** = *p* < 0.001. RPE = retinal pigment epithelium; Seg. = photoreceptor inner and outer segments; ONL = outer nuclear layer; OPL = outer plexiform layer; INL = inner nuclear layer; IPL = inner plexiform layer; GCL = ganglion cell layer; scale bar = 50 µm.

The PGC-1α expression pattern seen in in vitro retinal explant cultures was similar to that seen in in vivo retina (compare Figure 1C with Figure 3A). Quantitative immunofluorescence analysis showed that in ZLN005-treated *rd1* retinal explants, the fluorescence intensity of PGC-1α in the photoreceptor segments, IPL, and GCL was significantly stronger than that in the untreated *rd1* explants (Figure 3A, B), indicating that ZLN005 effectively upregulated PGC-1α expression.

Following ZLN005 treatment, PPARα fluorescence intensity was also significantly increased in the IPL and GCL (Figure 3A, B), suggesting that PGC-1α promoted PPARα expression. Notably, the expression pattern of PPARγ in the explants differed from that observed in the in vivo retina (Figure 1C; Figure 3A), such that PPARγ expression did not show significant changes in the segment layer but increased in the ONL, OPL, and INL (Figure 3A, B). In general, ZLN005 treatment significantly enhanced the expression of PGC-1α and upregulated the expression of PPARα and PPARγ, confirming PPARs/PGC-1α functional assembly in the retina.

### PPARs and PGC-1α activation modulate PARP activity

PGC-1α activity is tightly controlled via sirtuin-1-mediated deacetylation and we therefore investigated if selective activation of sirtuin-1 using resveratrol could restore its protective function^27^. However, in retinal explant cultures resveratrol treatment alone was insufficient to mitigate photoreceptor degeneration (Figure S3A). This lack of effect might have been due to the circumstance that NAD^+^, *i.e.* the substrate for sirtuin activity, was depleted by (excessive) PARP activity ^28^. We therefore utilized the specific inhibitor PJ34 to suppress PARP activity and to reduce NAD^+^ consumption. In line with previous data^29^, PJ34 treatment alone already significantly reduced photoreceptor cell death (Figure S3B). The combination of sirtuin-1 activation with resveratrol and PARP inhibition with PJ34 led to a further, statistically significant decrease in the number of ONL TUNEL-positive cells (Figure S3B). This data suggested that PARP, possibly via its use of NAD^+^, had an important effect on PPAR activity.

To further explore the link between PGC-1α signaling and PARP activity, *in situ* PARP activity assays and immunostaining for PAR were performed (Figure 4A-D; Figure S2A-D; Table S2 A, B). PARP activity and PAR-positive cells were lower in *wt* retinas compared with *rd1,* as shown previously^29^. However, ZLN005 treatment significantly reduced PARP activity and PAR generation in *rd1* ONL. The utilization of GW735 and Inolitazone also reduced PARP and PAR accumulation. In contrast, activation of PPARβ/δ using GW516 neither reduced PARP activity nor did it suppress PAR production. These results indicate that PGC-1α, PPARα, and PPARγ activation contribute to photoreceptor survival, at least in part, by suppressing PARP activity and its downstream effects. Taken together, this data suggests the existence of a feedback loop between the activities of PARP, sirtuin, PPAR, and PGC-1 in *rd1* photoreceptors.

**Figure 4.**
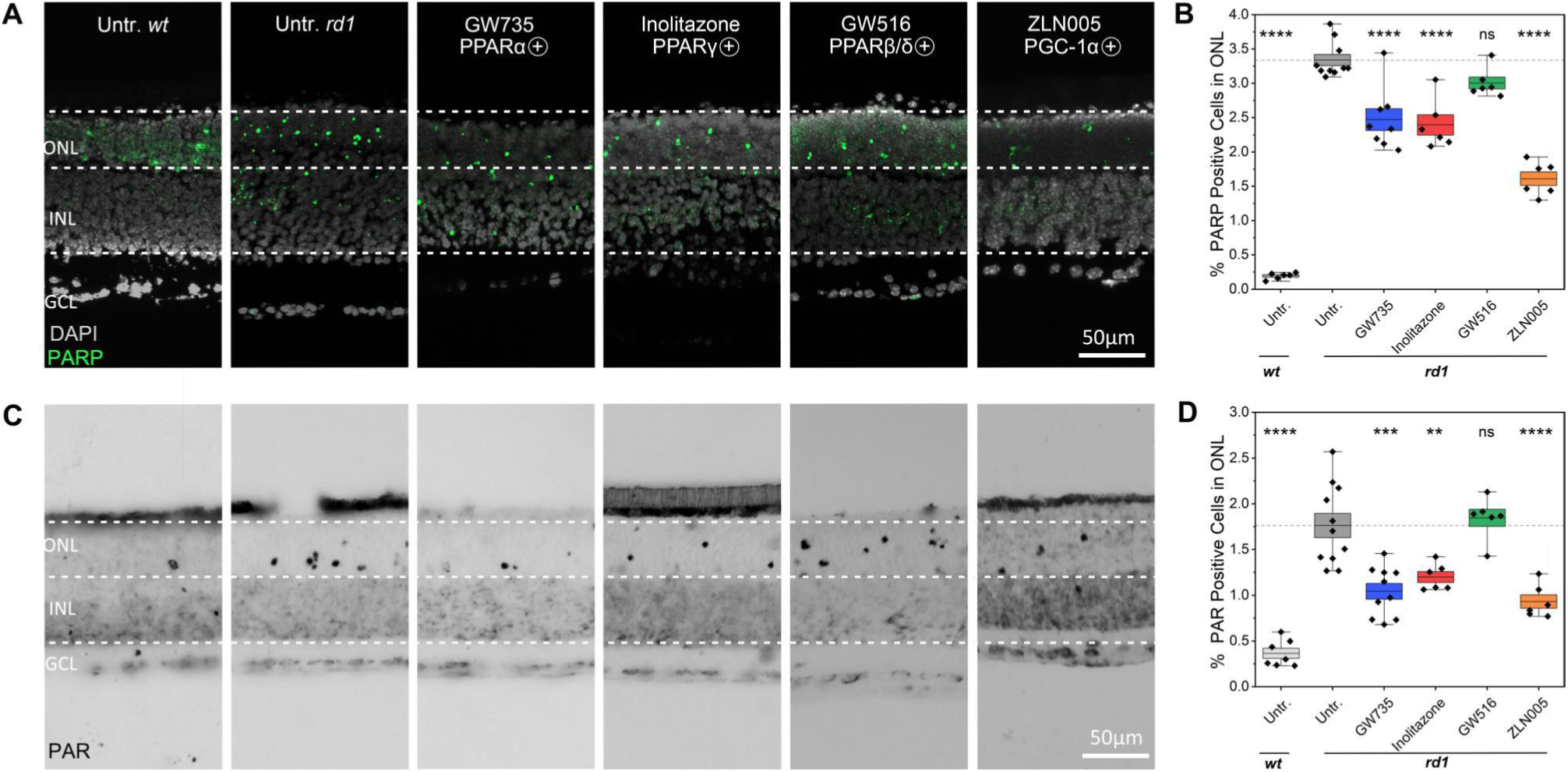
GW735, Inolitazone and ZL005 affect PARP activity and PAR generation. **A**) PARP activity assay (green) was performed in *rd1* and wild-type (*wt*) retinal explant cultures, with DAPI (grey) as nuclear counterstain. Untreated (Untr.) *wt* and *rd1* retina were compared to *rd1* retina treated with PPARα agonist GW735, PPARγ agonist Inolitazone, PPARβ/δ agonist GW516 and PGC-1α activator ZLN005. **B**) Box and whisker plots showing the percentage of PARP activity positive cells in the outer nuclear layer (ONL). The dashed line indicates *rd1* untr. situation, data points below this threshold indicate protective effects, data points above suggest destructive effects. **C**) PAR staining (black) was performed in *wt* and *rd1* retinal explant cultures. **D**) Untreated *wt* and *rd1* retina were compared to drug-treated retina as in B. Statistical testing: Two-way ANOVA and Dunnett’s multiple comparisons test. Error bars represent SD; significance levels: ns = *p* > 0.05; * = *p* < 0.05; ** = *p* < 0.01; *** = *p* < 0.001; **** = *p* < 0.0001. INL = inner nuclear layer, GCL = ganglion cell layer; scale bar = 50 µm.

### PPARγ and PGC-1α activation regulates calpain activity

Photoreceptor degeneration is also associated with calpain-mediated proteolysis and cellular damage. To investigate calpain activity, we first employed a general *in situ* calpain activity assay. While calpain activity was relatively low in *wt* compared to the *rd1* situation, treatment with Inolitazone and ZLN005 significantly reduced, while GW735 and GW516 did not affect, overall calpain activity (Figure 5A, D; Figure S2A-D; Table S3A).

**Figure 5.**
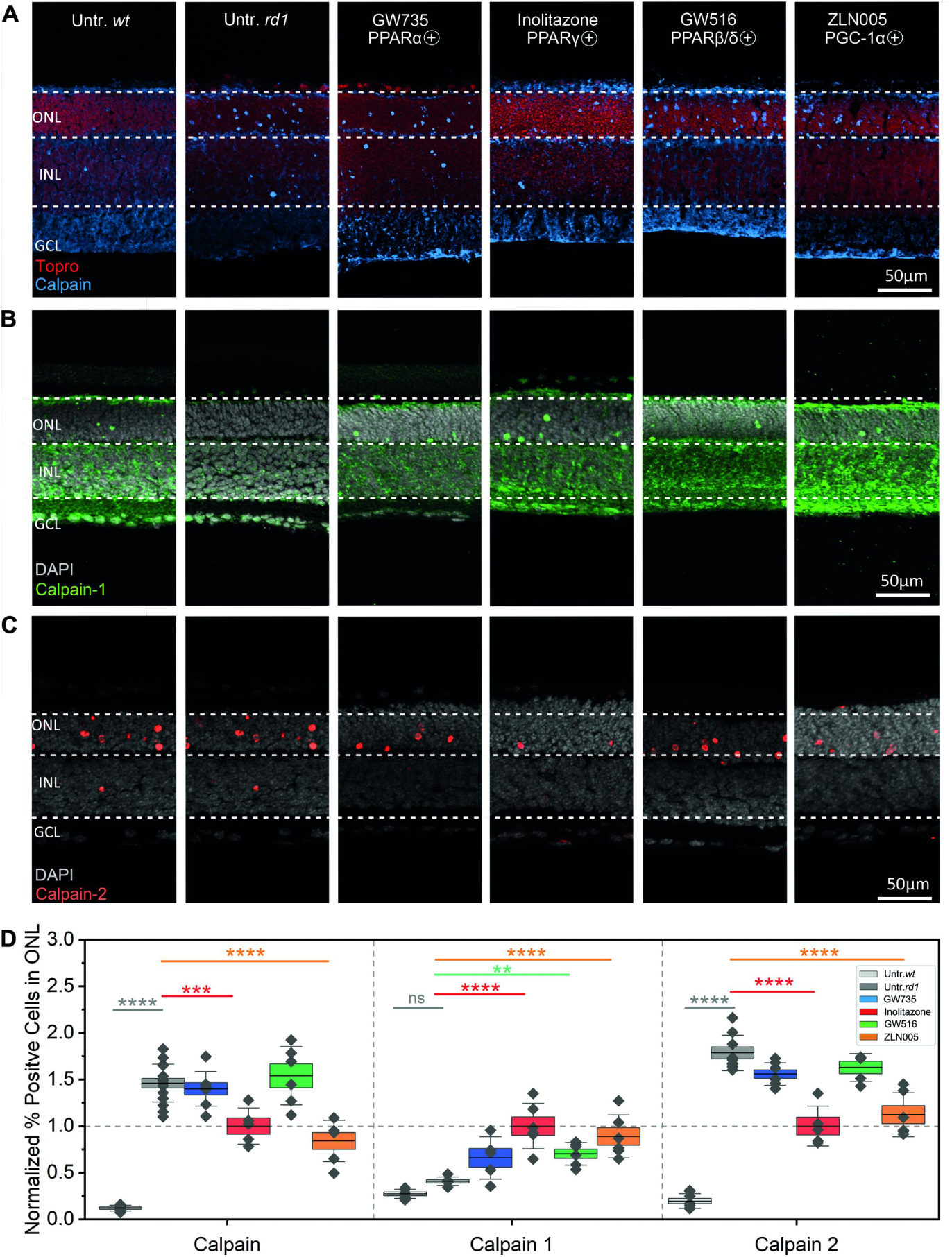
PPARs and PGC-1α differentially regulate calpain activity. **A**) Calpain activity assay (blue) in wild-type (*wt*) and *rd1* retinal explant cultures, nuclear counterstain with ToPro3 (red). Untreated (Untr.) *wt* and *rd1* retina were compared to *rd1* retina treated with PPARα agonist GW735, PPARγ agonist Inolitazone, PPARβ/δ agonist GW516 and PGC-1α agonist ZLN005. **B**) Activated calpain-1 immunostaining (green) in *wt* and *rd1* retinal explant cultures with DAPI (grey) as nuclear counterstain. **C**) Activated calpain-2 immunostaining (red) in *wt* and *rd1* retinal explant cultures with DAPI (grey) as nuclear counterstain. **D**) Box and whisker plot showing normalized percent calpain activity positive cells and displaying calpain-1, −2 activation in outer nuclear layer (ONL). Data was normalized to Inolitazone treatment (grey dashed line), which showed the strongest effect on calpain activity, according to the formula:χ_scaled_=χ∕χ_Inolitazone_, here, is the positive cells for the indicated condition, and _Inolitazone_, is the mean of Inolitazone treatment. Statistical comparisons were then performed using Untr. *rd1* as control to determine significant differences among treatments. Statistical testing: Two-way ANOVA and Dunnett’s multiple comparisons test. Error bars represent SD; significance levels: ns= *p* > 0.05; 0.01;** = *p* < 0.01; *** = *p* < 0.001; **** = *p* < 0.0001. INL = inner nuclear layer, GCL = ganglion cell layer; scale bar = 50 µm.

The two major calpain isoforms involved in neurodegenerative processes are calpain-1 and calpain-2, where calpain-1 activity is seen as mostly neuroprotective while activity of calpain-2 is likely destructive^30^. To study these two isoforms, we used immunolabelling for activated calpain-1 and calpain-2 and quantified the numbers of positive cells in the ONL (Figure 5B - D; Figure S2; Table S3B). The extent of calpain-1 activation was similar between *rd1* and *wt* retinas. Calpain-2 activation, however, was high in *rd1* retina when compared with *wt* retina. Inolitazone and ZLN005 significantly increased calpain-1 activation, while reducing calpain-2 activation. PPARβ/δ activation only increased the number of calpain-1-positive cells in the *rd1* ONL without inhibiting calpain-2 activation. There was no significant effect on the percentages of activated calpain-1 or calpain-2 positive cells in the *rd1* ONL after treatment with GW735.

## Discussion

In the CNS, PPARs play an important role in regulating cellular metabolism, as well as cell death and survival. Here, we show that PGC-1α and PPAR-signaling cooperate to maintain retinal homeostasis and photoreceptor cell survival in the context of inherited retinal degeneration (Figure 6). The protective effects of PGC-1α and PPAR activation are likely subject to positive regulation from sirtuin activity, while PARP activity may provide negative feedback. Moreover, the excessive activation of calpain-type proteases, notably calpain-2, appears to be situated down-stream of PGC-1α and PPAR-signaling.

**Figure 6.**
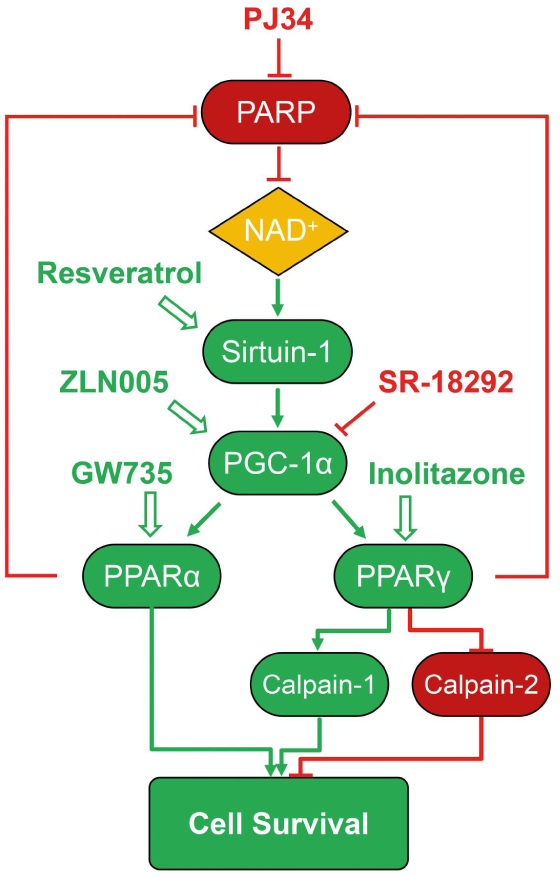
PGC-1α and PPAR retinal pro-survival signaling and experimental interventions. The PARP enzyme consumes NAD^+^, a substrate required for the function of sirtuin-1. Sirtuin-1 mediated deacetylation activates PGC-1α, which in turn functions as a coactivator for the nuclear receptors PPARα and PPARγ. This synergistic interaction drives transcriptional programs that support cell survival. While PPARα promotes survival independent of calpain activity, PPARγ signaling involves increased activation of the protective calpain-1 and decreased activation of the destructive calpain-2 isoform. This signaling network can be modulated by a variety of specific inhibitors and activators, potentially improving cell survival. While PARP can be inhibited by PJ34, resveratrol enhances sirtuin-1 activity as long as sufficient NAD^+^ is available. ZLN005 is described as a direct activator of PGC-1α, leading to its upregulation and activation. Conversely, SR-18292 inhibits PGC-1α. PPARα and PPARγ can also be directly targeted by the specific pharmacological agonists GW735 and Inolitazone, respectively. Red color indicates inhibition/downregulation, while green indicates activation/upregulation.

### PPARα-PPARγ-, and PGC-1α-activity contribute to photoreceptor survival

A general target of PPAR-signaling is lipid metabolism, which led us to the hypothesis that PPAR-associated abnormalities in lipid metabolism may play an important role in the pathogenesis of RP^31^. In particular, PPARα was found to be important for retinal fatty acid β-oxidation (FAO) and photoreceptor survival^32^. In *rd1* mouse retina, a downregulation of PPARα parallels the onset and progression of photoreceptor degeneration, suggesting a potential relationship between PPARα and retinal degeneration. The protective effect of the highly selective PPARα agonist GW735 may be mediated through enhancement of FAO, potentially improving energy supply to photoreceptors^33^.

The loss of PPARγ impairs fatty acid (FA) metabolism and leads to retinal abnormalities^32, 34^. While in *rd1* retina PPARγ gene expression was not significantly altered in the initial phases of retinal degeneration, treatment with the PPARγ agonist Inolitazone significantly increased photoreceptor survival. As PPARγ activation regulates FA transport^35^ and shift lipids from free fatty acid (FFA) to triglyceride (TG) storage^36^, Inolitazone may have protected photoreceptors via a decrease of lipid peroxidation through accelerated FFA clearance and optimized lipid distribution^37^.

In contrast, the transcriptomic data for PPARβ/δ show downregulated expression only in late stages of *rd1* rod degeneration. This indicates that PPARβ/δ changes may not be a primary driver of *rd1* cell death but rather a consequence of disease progression. Consistent with this idea, the PPARβ/δ agonist GW516 did not alleviate photoreceptor loss in *rd1* retina at a concentration that already induced toxicity in the *wt* control. This concept is in accordance with previous work that found widespread PPARβ/δ expression in the retina, but merely a limited role in maintaining normal retinal function^38^. Further, our findings confirm that PPARβ/δ activation does not rescue photoreceptors, with the implication that PPARβ/δ may not be suitable as a target for therapeutic intervention in RP.

PGC-1α is also recognized to play a key role in regulating lipid metabolism^39^, and its loss in retinal pigment epithelium (RPE) links abnormal lipid metabolism and retinal degeneration to PPAR control: Inhibition of PGC-1α downregulates PPARs expression, accompanied by reduced FAO, impaired FA transport, decreased triglyceride synthesis, and increased lipid peroxidation control^40^. We confirm an essential role of PGC-1α for photoreceptor viability in that its activation with ZLN005 enhances photoreceptor survival, while its inhibition with SR-18292 accelerates photoreceptor degeneration. Overall, our data thus reinforce the hypothesis that PPARα and PPARγ maintain lipid homeostasis through a mechanism coordinated by PGC1α. Another important mechanism concerns the role of PGC-1α to drive mitochondrial biogenesis, including in RP^41^, and that PPARs can also cooperate with PGC-1α to advance this process^42, 43^. Hence, PGC-1α may cooperate with PPARs to correct lipid metabolic disturbances, indirectly boosting mitochondrial function.

### Bidirectional control of PPAR-signaling via PARP activity

Full activation of PGC-1α is achieved by sirtuin-1 mediated deacetylation^44^ and thus the sirtuin activator resveratrol may exert a neuroprotective function^45, 46^. Yet, in *rd1* mice, resveratrol failed to prevent photoreceptor degeneration consistent with a previous study on the N-methyl-N-nitrosourea (MNU) model for RP, in which resveratrol also did not prevent the decline of retinal function^47^. To be active, sirtuin-1 requires NAD^+^ as substrate, which on the other hand is also the substrate for PARP activity. As the PARP overactivation observed in *rd1* photoreceptors likely depletes NAD^+^, it would limit sirtuin-1 activity. This interpretation is consistent with previous studies^48^ and supported by the results obtained from the combined treatment using resveratrol to activate sirtuin-1 and PJ34 to block PARP, which significantly decreased ONL cell death. Taken together, this implies that PARP is an upstream regulator of PGC-1α/PPARs-signaling.

On the other hand, treatment with PGC-1α, PPARα and PPARγ activators reduced photoreceptor PARP activity and PAR accumulation, indicating that PARP is also a downstream effector. It has been reported that PGC-1α, PPARα, and PPARγ reduce superoxide production and activate the expression of antioxidant enzymes^49–53^. Thus, PGC-1α/PPAR-signaling may reduce PARP activity possibly by maintaining redox homeostasis and limiting oxidative damage to DNA. Overall, our data support the concept of a PARP–sirtuin-1–PGC-1α–PPAR–PARP bidirectional feedback loop that may couple a deficient energy metabolism to photoreceptor degeneration (Figure 6).

### PPARγ and PGC-1α regulate calpain proteolytic activity

Regarding calpains, we found that the activation of PGC-1α and PPARγ produced a marked downregulation of overall calpain activity and specifically a reduction of calpain-2 activation. Since calpain-2 activation requires high intracellular Ca^2+^-levels, we hypothesize a role of PPARγ in the maintenance of cellular Ca^2+^ homeostasis. This is supported by the fact that PPARγ agonists are known to maintain mitochondrial Na^+^/Ca^2+^ exchanger levels allowing mitochondria to buffer intracellular Ca^2+54^. Furthermore, PGC-1α can prevent mitochondrial Ca^2+^ overload by regulating expression of the mitochondrial calcium uniporter complex^55, 56^. It is thus plausible to think that PGC-1α together with PPARγ regulates intracellular Ca^2+^-levels and indirectly calpain activation. Eventually, this may lead to an excessive proteolysis, which, without further control, would promote photoreceptor cell death. On the other hand, both PGC-1α and PPARγ activation resulted in increased activation of calpain-1, an isoform often associated with neuroprotection^30^, confirming a recent study on the mechanisms of *rd1* photoreceptor degeneration^57^.

In contrast, PPARβ/δ activation had no effect on photoreceptor calpain activity, a result which corresponds to its lack of effect on *rd1* ONL cell death, and possibly explained by its localization to the retinal vasculature.

Perhaps the most interesting observation in this context is that PPARα activation, while reducing ONL cell death in the *rd1* model, did not significantly affect calpain activity. This indicates that the neuroprotection accomplished by PPARα is largely independent of Ca^2+^ homeostasis regulation. Altogether, these observations provide a rationale for a combined treatment with PPARα and PPARγ agonists to exploit the full therapeutic power provided by different and independent neuroprotective mechanisms.

### Cooperation and differentiation of PPARα, PPARγ, and PGC-1α

The different localizations of PPARα, PPARγ, and PGC-1α in the retina suggest that their roles are cell-specific, which also affects their contributions during photoreceptor degeneration. Both PPARα and PPARγ were expressed in the extremely lipid-membrane rich photoreceptor OS, suggesting a function in the regulation of lipid metabolism and the maintenance of OS structure. PPARα was additionally expressed across essentially the entire retina, implying that PPARα may not only help to maintain metabolic homeostasis within photoreceptors, but also in other retinal cell types, including perhaps Müller glia cells^58, 59^. PGC-1α featured a broader expression pattern in the retina, widely distributed in the ONL, IPL, and INL, indicating that it may be involved in regulating lipid metabolism, energy supply, mitochondrial homeostasis, and antioxidant responses across multiple retinal layers. Although PPARα, PPARγ, and PGC-1α possibly differ in function in different retinal cell types, their regulatory mechanisms partially overlap, particularly in regulating PARP activity ^60, 61^. Nevertheless, the differential effects of PPARα and PPARγ agonists on calpain activity highlight distinct downstream regulatory mechanisms of PPARα, PPARγ, and PGC-1α in RP.

## Conclusion

The key finding of this work is that activation of PPARα, PPARγ, and PGC-1α is able to alleviate photoreceptor degeneration in the retina of a murine model of RP. These factors may synergistically interact to improve photoreceptor viability through several pathways, including fatty acid oxidation, maintenance of mitochondrial homeostasis, and inhibition of calpain and PARP overactivation. Moreover, their selective regulation reveals distinct neuroprotective mechanisms for different PPAR subtypes and PGC-1α, providing a foundation for future intervention strategies in degenerative retinal diseases improved by addressing different targets in a combined approach. The exploration and development of such a novel therapeutic strategy for RP will certainly profit from a further validation via transcriptional, metabolic, and mitochondrial analyses to clarify causal mechanisms and map the complete PGC-1α/PPAR signaling network.

## Materials and methods

### Animals

Wild-type (*wt*) C3H/HeA *Pde6b^+/+^* and congenic C3H/HeA *Pde6b^rd1/rd1^* mice (*rd1*) were used^62^ irrespective of sex. All efforts were made to minimize the number of animals used and their suffering. Mice were housed in the specified-pathogen-free (SPF) housing facility at the Tübingen Institute for Ophthalmic Research, under standard white cycling light, with free access to food and water.

### Analysis of RNA-seq data

The mRNA expression comparison between *wt* and *rd1* mouse employed the GSE62020 dataset downloaded from the Gene Expression Omnibus (GEO) database (https://www.ncbi.nlm.nih.gov/geo; information retrieved in July 2025). This dataset was previously reporte^23^. To characterize *rd1* data, analysis was performed using R software (version 4.5.1) and the limma package (version 3.64.3) for normalization and differential expression analysis (fold change > 1.2, p < 0.05)^63^.

### Organotypic retinal explant cultures

Retinas were used to generate explants following the standard protocol as previously described^64^. Mice were decapitated at postnatal day 5 (P5), the heads cleaned with 70% ethanol and the eyes removed under aseptic conditions. The explants were cultured on a polycarbonate membrane (Order No.: 83.3930.040; 0.4 µm TC-inserts; Sarstedt, Nümbrecht, Germany) with complete R16 medium (Order No: 07491252A, Gibco, Paisley, UK) including supplements^64^. On the day of culture, retinal explants from left and right eyes of each mouse were randomly assigned to different experimental groups, subsequent processing was identical across groups. No blinding was used for sample allocation. The medium was exchanged every two days with treatment added at P7 and P9. Cultures were treated with 0.04 µM GW590735 (HY-106278, MedChemExpress, Sollentuna, Sweden), 0.8 µM Inolitazone dihydrochloride (HY-14792B; MedChemExpress), 1 µM GW501516 (HY-10838; MedChemExpress), 0.5 µM ZLN005 (HY-17538; MedChemExpress), 20 µM SR-18292 (HY-101491; MedChemExpress), 20 µM resveratrol (501-36-0; Sigma, St.Louis, MO, USA), 6 µM PJ34 (Alexis Biochemicals, Lausen, Switzerland). All compounds were dissolved in DMSO at a final medium concentration of no more than 0.1% DMSO. Culturing was stopped at P11 by either fixation with 4% paraformaldehyde (PFA) or without fixation and direct freezing in liquid N_2_ for all treatment conditions. Explants were embedded in Tissue-Tek O.C.T. compound (Sakura Finetek Europe B.V., Alphen aan den Rijn, The Netherlands), then cut into 14 µm sections using a cryostat (CryoStar NX50 OVP; Thermo Fisher Scientific, Runcorn, UK), after which they were dried and stored at −20°C for future use.

### TUNEL Assay

A terminal deoxynucleotidyl transferase dUTP nick end labeling (TUNEL) assay kit (Roche Diagnostics, Mannheim, Germany) was used to label dying cells in retinal tissue sections. Fixed sections were rehydrated with 0.1 M phosphate-buffered saline (PBS) at room temperature (RT), while unfixed sections were first fixed in 4% PFA for 10 min, then washed with PBS. Subsequent steps were identical for both fixed and unfixed sections. Sections were incubated in Tris-buffer saline (TBS; 1 µL enzyme/mL) with proteinase K (1.5 µg/µL) at 37 °C for 5 min to inactivate nucleases, followed by treatment with 70:30 ethanol-acetic acid mixture at −20 °C for 5 min and three washes in TBS. Sections were then incubated with blocking solution (10% normal goat serum (NGS), 1% bovine serum albumin (BSA), and 1% fish gelatin in PBS with 0.3% Triton X-100 for 1 hour at RT, followed by overnight incubation with TUNEL staining solution at 4 °C. Finally, sections were washed three times with PBS (5 min each), mounted in Vectashield with DAPI (Vector Laboratories Inc., Burlingame, CA, USA), and imaged by microscopy.

### Calpain activity assay

Unfixed retinal tissue sections were rehydrated in calpain reaction buffer (CRB) (5.96 g HEPES, 4.85 g KCl, 0.47 g MgCl2, and 0.22 g CaCl2 in 100 mL ddH2O; pH 7.2) with 2 mM dithiothreitol (DTT) for 15 min. Then, sections were incubated with 25 µM tBOC-Leu-Met-CMAC (A6520; Thermo Fisher Scientific, OR, USA) for 3 h at 37 °C. This was followed by two washes with PBS and 12 min incubation with ToPro3 (1:1000 in PBS; T3605, Invitrogen, Carlsbad, USA). Afterwards, sections were mounted using Vectashield without DAPI (Vector Laboratories) for immediate microscopic visualization.

### PARP activity

The *in situ* PARP activity assay was performed as published previously^65^. In brief, unfixed retinal tissue sections were rehydrated in PBS for 10 min, then incubated at 37 °C for 3 h in a reaction mixture (10 mM MgCl_2_, 1 mM DTT, and 50 µM 6-Fluo-10-NAD^+^ (Cat. Nr.: N 023; Biolog, Bremen, Germany) in 100 mM Tris buffer with 0.2% Triton X100, pH 8.0). After three 5-min washes in PBS, sections were mounted in Vectashield with DAPI (Vector Laboratories).

### PAR-DAB staining

3,3′-diaminobenzidine (DAB) staining was performed on fixed sections to detect PAR generation. To quench endogenous peroxidase activity, sections were incubated in 40% Methanol and 10% H_2_O_2_ in PBS with 0.3% Triton X-100 (PBST) for 20 min. Sections were further incubated with 10% NGS in PBST for 30 min, followed by anti-PAR antibody (1:200; Enzo Life Sciences, Farmingdale, New-York) incubation, overnight, at 4 °C. After three 10 min washes in PBS, sections were incubated with biotinylated secondary antibody (1:150, Vector in 5% NGS in PBST) for 1 h. This was followed by staining with the Vector ABC Kit (Vector Laboratories, solution A and solution B in PBS, 1:150 each) for 1 h. DAB staining solution (0.05 mg/mL NH_4_Cl, 200 mg/mL glucose, 0.8 mg/mL nickel ammonium sulfate, 1 mg/mL DAB, and 0.1 vol. % glucose oxidase in phosphate buffer) was applied evenly, incubated for 2 min, and immediately rinsed with phosphate buffer to stop the reaction. Sections were mounted in Aquatex (Merck, Darmstadt, Germany).

### Immunohistochemistry

Fixed eyecups or retinal explant slides were rehydrated in PBS for 15 min at RT. Afterwards, sections were incubated with blocking solution (10% NGS, 1% BSA, 0.3% PBST) for 1 h. The primary antibodies (Table 1) were diluted in blocking solution and incubated overnight at 4 °C. Rinsing with PBS 3 times 10 min each was followed by incubation with secondary antibody (Table 1) for one hour. The sections were rinsed with PBS 3 times, 10 min each, and mounted with mounting medium with DAPI (Vector Laboratories).

**Table 1.** Primary and secondary antibodies used in the study, providers, and dilutions.

| Antibody | Provider, Cat. No. | Dilution |
| --- | --- | --- |
| Rabbit polyclonal anti-calpain-1 | Abcam, ab39170 | 1:100 |
| Rabbit polyclonal anti-calpain 2 | Abcam, ab39165 | 1:200 |
| Mouse monoclonal anti-glutamine synthetase (GS) | Abcam, MAB302 | 1:1000 |
| Rabbit polyclonal anti-PPAR alpha | Novus, NB600-636 | 1:100 |
| Rabbit polyclonal anti-PPAR gamma | Proteintech, 16643-1-AP | 1:200 |
| Mouse monoclonal anti-PPAR delta | Immunoway, YM3601 | 1:100 |
| Mouse monoclonal anti-PGC-1 alpha | Proteintech, 66369-1-Ig | 1:200 |
| Peanut agglutinin (PNA) | Vector laboratories, FL-1071 | 1:500 |
| Goat-anti-rabbit AlexaFluor568 (AF568) | Molecular Probes, A11036 | 1:350 |
| Goat-anti-rabbit AlexaFluor488 (AF488) | Molecular Probes, A11034 | 1:350 |
| Goat-anti-mouse AlexaFluor568 (AF568) | Molecular Probes, A11031 | 1:350 |

### Microscopy and image analysis

Images were captured using a Zeiss Imager Z.2 fluorescence microscope equipped with ApoTome2, an Axiocam 506 mono camera, and an HXP-120V fluorescent lamp (Carl Zeiss Microscopy, Oberkochen, Germany). The excitation (λExc)/emission (λEm) characteristics of the filter sets used for the different fluorophores were as follows (in nm): DAPI (λExc = 369 nm, λEm = 465 nm), AF488 (λExc = 490 nm, λEm = 525 nm), AF568 (λExc = 578 nm, λEm = 602 nm), and ToPro (λExc = 642 nm, λEm = 661 nm). Images were captured at 20 × magnification using the Z-stack mode of Zen 2.3 blue edition software. Sections of 14 μm thickness were analyzed using 15 Apotome Z-planes. Data were obtained from *ex vivo* retinas or explants derived from at least three animals for each condition.

For quantification, images were captured at six random locations per retinal section. The percentage of positive cells was determined as follows: The average area occupied by a photoreceptor cell (*i.e*., cell size) was determined by counting the number of DAPI-stained nuclei in six different rectangular areas of the retina. The total number of photoreceptor cells was estimated by dividing the given ONL area by this average cell size. The number of positively labeled cells (TUNEL, PARP, PAR, calpain, calpain-1, calpain-2) in the ONL was counted manually. The percentage of positive cells was then calculated by dividing the number of positive cells by the total number of ONL cells.

The signal intensity on retinal sections was quantified by Zen software version 2.3 (Zeiss). DAPI staining was used to identify the location of different retinal layers, and fluorescence intensity was manually recorded for each layer. For each count, fluorescence intensity was assessed at nine distinct positions. The values of all sections from the same animal were averaged before further analysis.

### Statistical analysis and software use

Minimum animal and sample numbers were determined by an a priori power analysis based on effect sizes from previous explant studies^29, 57^, performed using StatistikGuru (v1.96; https://www.statistikguru.de; accessed Oct. 2025). Animals were not allocated to experimental groups prior to sacrifice, and explants from both eyes of all animals were included without exclusion. Two-way comparisons were analyzed using Student’s *t-*test. Multiple comparisons were made using a one-way analysis of variance (ANOVA) test or two-way ANOVA with Dunnett’s multiple comparisons test. All tests were two-sided, normality was checked by Shapiro-Wilk, and variance homogeneity was evaluated using the Brown-Forsythe test. GraphPad Prism 10.6.1 software (GraphPad Software, San Diego, CA, USA) was used for statistical analysis and data visualization. Box-and-whisker plots were generated in OriginPro 2024b (OriginLab, Northampton, MA, USA). Levels of significance were as follows: ns, *p* > 0.05; *, *p* < 0.05; **, *p* < 0.01; ***, *p* < 0.001; ****, *p* < 0.0001. The figures were prepared using Adobe Photoshop 2020 (San Jose, CA, USA). Analysis of transcriptome datasets was carried out using Cytoscape 3.10.4 software (Cytoscape consortium) with the Molecular Complex Detection (MCODE) plugin^24^ for module clustering and the cytoHubba plugin for hub gene identification and ranking^25^.

## Acknowledgments

The authors would like to thank Norman Rieger (from Institute for Ophthalmic Research, Eberhard-Karls-Universität Tübingen) for excellent technical assistance.

## Author contributions

Conceptualization, L.W., J.Y., and F.P.-D.; methodology, L.W., and F.P.-D.; software, L.W., J.Y., and QL.Y; validation, L.W.; formal analysis, L.W., J.Y., and QL.Y.; investigation, L.W., J.Y., KW.J., ZL.H., and F.P.-D.; data curation, L.W., J.S-P.; writing-original draft preparation, L.W.; writing-review and editing, F.P-D. and M.S.; visualization, L.W., J.Y., and QL.Y.; supervision, F.P-D.; project administration, F.P-D.; funding acquisition, F.P-D. All authors have read and agreed to the published version of the manuscript.

## Ethical approval and consent to participate

The animal study protocols compliant with the German law on animal protection were reviewed and approved by the institutional animal welfare committee of the University of Tübingen (Registration Nos: AK02/24M, AK05/22M) and were following the association for research in vision and ophthalmology (ARVO) statement for the use of animals in vision research.

## Data availability

All data generated or analysed during this study are included in this published article and its Supplementary Materials Files.

