## Supplemental figures and tables for "PGC-1α and PPARs cooperatively mediate photoreceptor neuroprotection in *rd1* mouse inherited retinal degeneration"

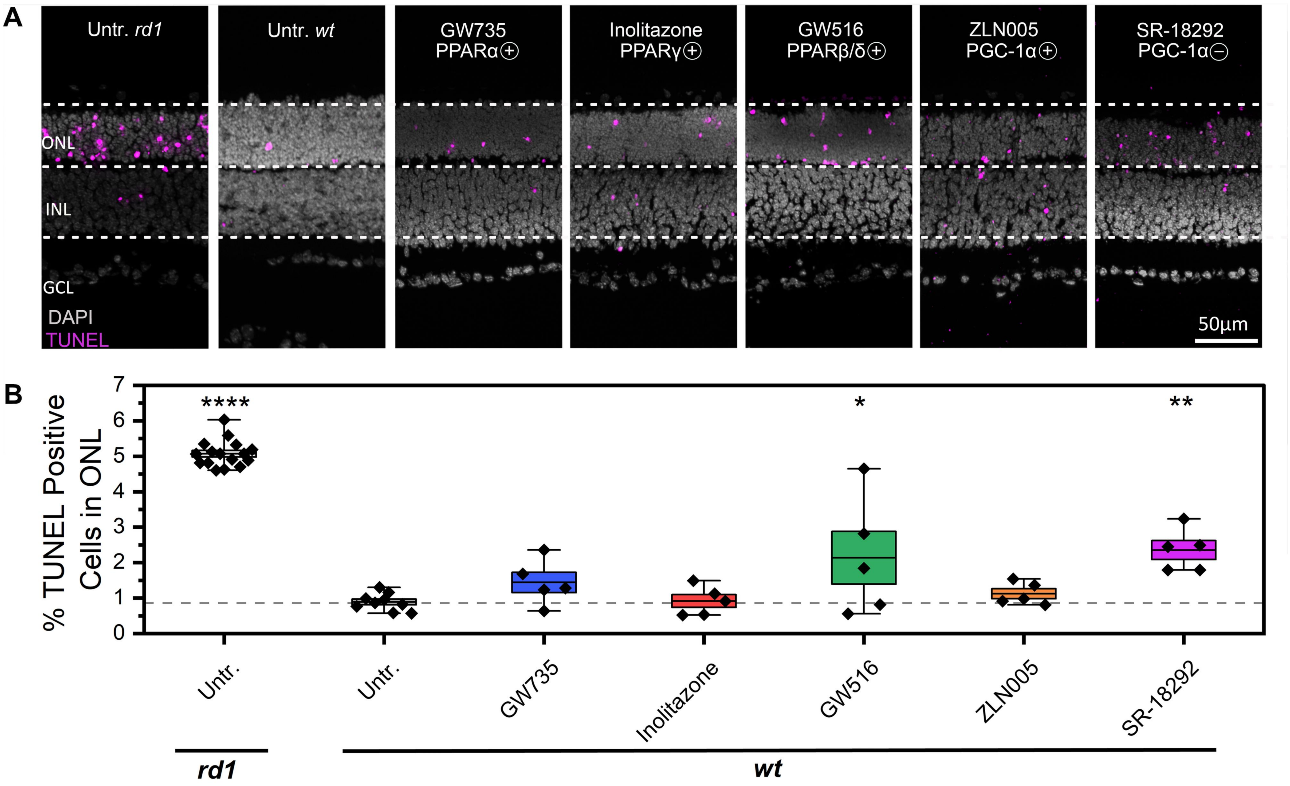


**Figure S1. GW516 and SR-18292 increase wild-type photoreceptor cell death. A)** TUNEL positive cells (magenta) in wild-type (*wt*) and *rd1* retinal cultures. *wt* retina was left either untreated (Untr.) or treated with PPARα agonist GW735, PPARγ agonist Inolitazone, PPARβ/δ agonist GW516, PGC-1α activator ZLN005 and inhibitor SR-18292. **B**) Box and whisker plot showing percentages of TUNEL positive cells in the outer nuclear layer (ONL). The grey dashed line designates *wt* untr. situation, data points below this threshold indicate protective effects, data points above suggest destructive effects. Statistical testing: Two-way ANOVA with Dunnett's multiple comparisons test performed between *rd1* and *wt* explant cultures. Error bars represent SD; * = $p < 0.05$; ** = $p < 0.01$; **** = $p < 0.0001$. INL = inner nuclear layer, GCL = ganglion cell layer; scale bar = 50 µm

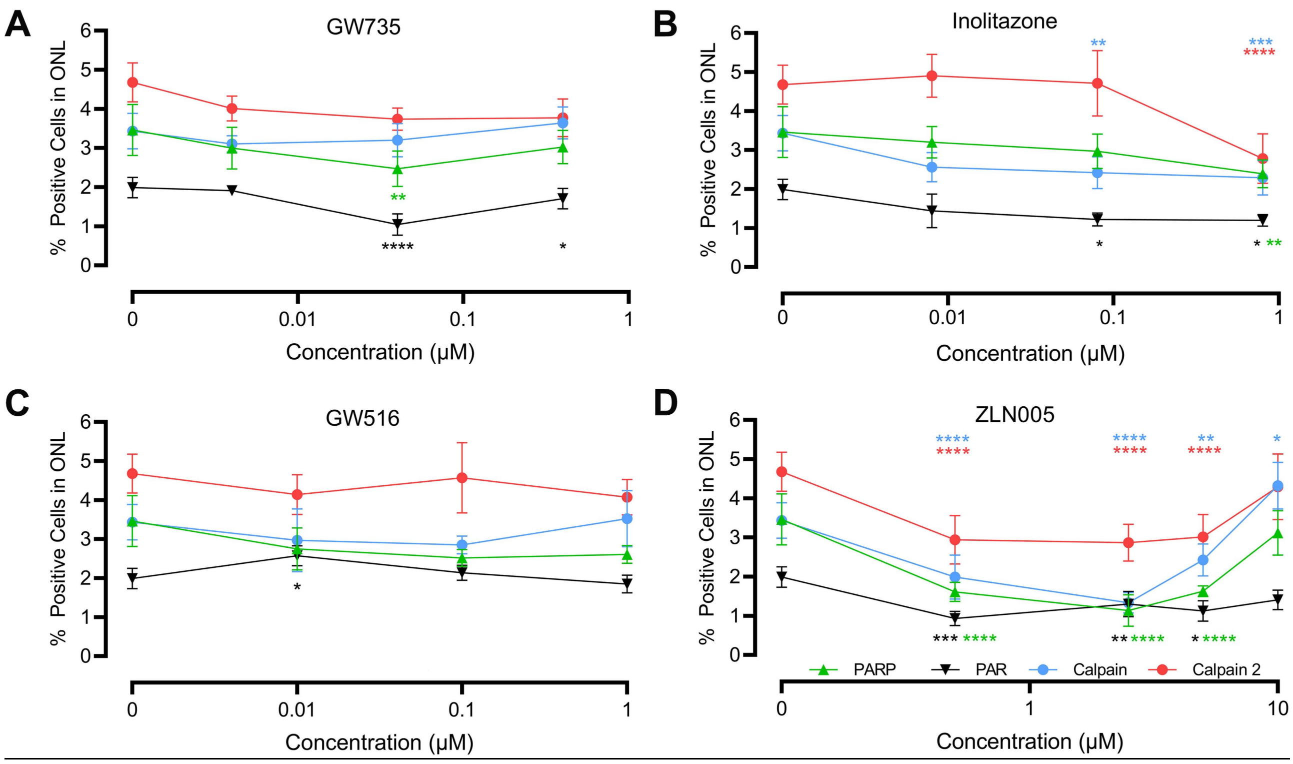


**Figure S2. Dose-response curves for GW735, Inolitazone, GW516, and ZLN005.** Markers assessed in the outer nuclear layer (ONL) of *rd1* retinal explant cultures were *in situ* PARP activity (green lines), accumulation of PAR (black), *in situ* calpain activity (blue), and activation of calpain-2 (red). **A**) In the ONL, at concentrations of 0.04 μM, the PPARα activator GW735 significantly decreased both the numbers of cells displaying PARP activity and PAR accumulation, at 0.4 μM it significantly reduced PARP activity. **B**) At concentrations of 0.8 μM, the PPARγ activator Inolitazone significantly reduced ONL PARP activity, PAR generation, calpain activity, and calpain 2 activation. 0.08 μM Inolitazone significantly reduced PAR generation and calpain activity. **C**) 0.01 μM of the PPAR β/δ activator GW516 significantly increased PAR production in the *rd1* ONL. **D**) The PGC-1α activator ZLN005, over a broad range of concentrations (0.5-5 μM), significantly reduced PARP activity, PAR generation, calpain activity, and calpain-2 activation in the ONL. Yet, at 10 μM, ZLN005 significantly increased calpain activity. Statistical significance was assessed using one-way ANOVA and Dunnett's multiple comparison test; significance levels: * = $p < 0.05$; ** = $p < 0.01$; *** = $p < 0.001$; **** = $p < 0.0001$; error bars represent SD.

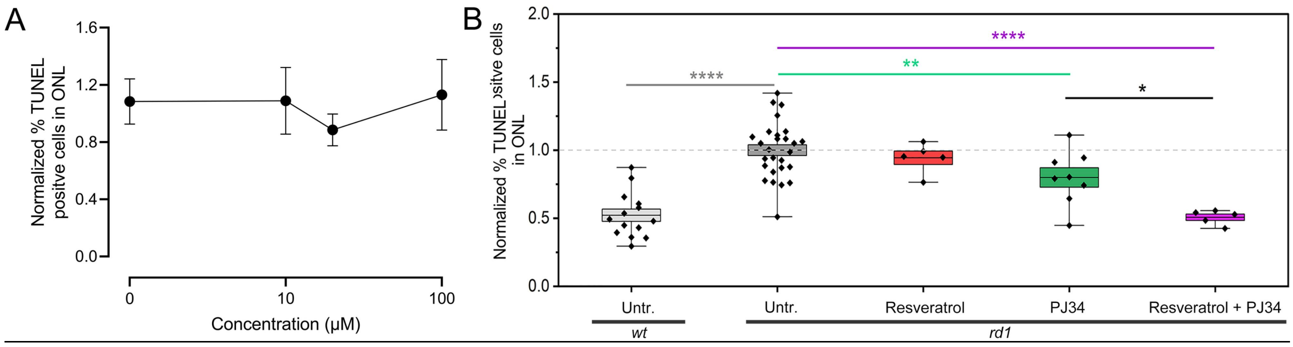


**Figure S3. Sirtuin-1 dependent photoreceptor protection requires PARP inhibition. A)** Dose-response curve for the sirtuin-1 agonist resveratrol (0 – 100 µM) in *rd1* mouse retinal explant cultures. Cell death in *rd1* outer nuclear layer (ONL) was assessed using the TUNEL assay, normalized to the untreated (Untr.) *rd1* situation. Resveratrol did not display significant *rd1* photoreceptor protection at any concentration tested. **B)** Combination treatment with sirtuin-1 agonist and PARP antagonist. The grey dashed line designates *rd1* untr. situation, data points below this threshold indicate protective effects, data points above suggest destructive effects. In the ONL, at 20 µM, resveratrol did not decrease the number of TUNEL positive cells, while 6 µM of the PARP inhibitor PJ34 significantly reduced the number of dying cells. However, 20 µM resveratrol combined with 6 µM PJ34 further reduced TUNEL positive cells in *rd1* ONL. Statistical significance was assessed using one-way ANOVA and Dunnett's multiple comparison test in A, two-way ANOVA with Dunnett's multiple comparisons test in B; significance levels: * = $p < 0.05$; ** = $p < 0.01$; **** = $p < 0.0001$; error bars represent SD. Data was normalized by linear scaling according to the formula: $\chi_{\text{scaled}} = \chi / \overline{\chi}_{rd1,\text{Untr.}}$, here, $\chi$ is the TUNEL positive cells for the indicated condition, and $\overline{\chi}_{rd1,\ \text{Untr.}}$, is the mean of *rd1* untreated controls.

**Supplemental Table 1. Quantification of TUNEL positive, dying cells in the outer nuclear layer (ONL).**

| A | | | Mean % positive cells in ONL | Standard deviation (SD) | Sample (n) | Dunnett's multiple comparisons test | Significance(*p*) |
|---|---|---|---|---|---|---|---|
| *rd1* | | *rd1* Untr. | 5.07 | ±0.38 | 16 | *rd1* Untr. vs *wt* Untr. | *P* < 0.0001 |
| | GW735 | 0.004 µM | 4.69 | ±0.82 | 9 | *rd1* Untr. vs 0.004 µM GW735 | *P* > 0.05 |
| | | 0.04µM | 3.84 | ±0.37 | 9 | *rd1* Untr. vs 0.04 µM GW735 | *P* < 0.01 |
| | | 0.4µM | 4.23 | ±0.90 | 9 | *rd1* Untr. vs 0.04 µM GW735 | *P* > 0.05 |
| | Inolitazone | 0.008 µM | 4.76 | ±0.81 | 12 | *rd1* Untr. vs 0.008 µM Inolitazone | *P* > 0.05 |
| | | 0.08 µM | 4.21 | ±0.57 | 12 | *rd1* Untr. vs 0.08 µM Inolitazone | *P* > 0.05 |
| | | 0.8 µM | 3.68 | ±0.54 | 12 | *rd1* Untr. vs 0.8 µM Inolitazone | *P* < 0.001 |
| | GW516 | 0.01 µM | 4.68 | ±1.27 | 11 | *rd1* Untr. vs 0.01 µM GW516 | *P* > 0.05 |
| | | 0.1 µM | 4.40 | ±0.92 | 11 | *rd1* Untr. vs 0.1 µM GW516 | *P* > 0.05 |
| | | 1 µM | 4.30 | ±0.60 | 10 | *rd1* Untr. vs 1 µM GW516 | *P* > 0.05 |
| | ZLN005 | 1 µM | 3.24 | ±0.42 | 10 | *rd1* Untr. vs 0.5 µM ZLN005 | *P* < 0.0001 |
| | | 2.5 µM | 3.24 | ±0.83 | 9 | *rd1* Untr. vs 2.5 µM ZLN005 | *P* < 0.0001 |
| | | 5 µM | 3.51 | ±0.49 | 7 | *rd1* Untr. vs 5 µM ZLN005 | *P* < 0.001 |
| | | 10 µM | 5.09 | ±0.93 | 5 | *rd1* Untr. vs 10 µM ZLN005 | *P* > 0.05 |
| | SR-18292 | 1 µM | 5.57 | ±0.47 | 6 | *rd1* Untr. vs 1 µM SR-18292 | *P* > 0.05 |
| | | 5 µM | 4.58 | ±0.15 | 6 | *rd1* Untr. vs 5 µM SR-18292 | *P* > 0.05 |
| | | 10 µM | 5.72 | ±1.25 | 6 | *rd1* Untr. vs 10 µM SR-18292 | *P* > 0.05 |
| | | 20 µM | 20.03 | ±3.00 | 3 | *rd1* Untr. vs 20 µM SR-18292 | *P* < 0.0001 |

| B | | Mean % positive cells in ONL | Standard deviation (SD) | Sample (n) | Dunnett's multiple comparisons test | Significance(*p*) |
|---|---|---|---|---|---|---|
| C3H | *wt* Untr. | 0.89 | ±0.24 | 9 | *wt* Untr. vs *rd1* Untr. | *P* < 0.0001 |
| | GW735 | 1.44 | ±0.63 | 5 | *wt* Untr. vs GW735 | *P* > 0.05 |
| | Inolitazone | 0.92 | ±0.41 | 5 | *wt* Untr. vs Inolitazone | *P* > 0.05 |
| | GW516 | 2.32 | ±1.67 | 5 | *wt* Untr. vs GW516 | *P* < 0.05 |
| | ZLN005 | 1.12 | ±0.32 | 5 | *wt* Untr. vs ZLN005 | *P* > 0.05 |
| | SR-18292 | 2.35 | ±0.60 | 5 | *wt* Untr. vs SR-18292 | *P* < 0.01 |

**Supplemental Table 2. Quantification of PARP activity and PAR positive cells in the outer nuclear layer (ONL).**

| A | | Mean % positive cells in ONL | Standard deviation (SD) | Sample (n) | Dunnett's multiple comparisons test | Significance(*p*) |
|---|---|---|---|---|---|---|
| PARP activity | *wt* Untr. | 0.19 | ±0.05 | 6 | *rd1* Untr. vs *wt* Untr. | *P* < 0.0001 |
| | *rd1* Untr. | 3.34 | ±0.26 | 10 | | |
| | GW735 | 2.47 | ±0.45 | 8 | *rd1* Untr. vs GW735 | *P* < 0.0001 |
| | Inolitazone | 2.40 | ±0.36 | 6 | *rd1* Untr. vs Inolitazone | *P* < 0.0001 |
| | GW516 | 3.01 | ±0.21 | 6 | *rd1* Untr. vs GW516 | *P* > 0.05 |
| | ZLN005 | 1.61 | ±0.23 | 6 | *rd1* Untr. vs ZLN005 | *P* < 0.0001 |

| B | | Mean % positive cells in ONL | Standard deviation (SD) | Sample (n) | Dunnett's multiple comparisons test | Significance(*p*) |
|---|---|---|---|---|---|---|
| PAR | *wt* Untr. | 0.37 | ±0.15 | 7 | *rd1* Untr. vs *wt* Untr. | *P* < 0.0001 |
| | *rd1* Untr. | 1.76 | ±0.44 | 11 | | |
| | GW735 | 1.04 | ±0.27 | 10 | *rd1* Untr. vs GW735 | *P* < 0.001 |
| | Inolitazone | 1.20 | ±0.15 | 6 | *rd1* Untr. vs Inolitazone | *P* < 0.05 |
| | GW516 | 1.85 | ±0.23 | 6 | *rd1* Untr. vs GW516 | *P* > 0.05 |
| | ZLN005 | 1.10 | ±0.57 | 6 | *rd1* Untr. vs ZLN005 | *P* < 0.01 |

**Supplemental Table 3. Quantification of calpain activity, calpain-1 activation, and calpain-2 activation in the outer nuclear layer (ONL).**

| A | | Mean % positive cells in ONL | Standard deviation (SD) | Sample (n) | Dunnett's multiple comparisons test | Significance(*p*) |
|---|---|---|---|---|---|---|
| Calpain activity | *wt* Untr. | 0.31 | ±0.12 | 7 | *rd1* Untr. vs *wt* Untr. | *P* < 0.0001 |
| | *rd1* Untr. | 3.34 | ±0.46 | 16 | | |
| | GW735 | 3.09 | ±0.28 | 8 | *rd1* Untr. vs GW735 | *P* > 0.05 |
| | Inolitazone | 2.29 | ±0.44 | 5 | *rd1* Untr. vs Inolitazone | *P* < 0.001 |
| | GW516 | 3.52 | ±0.72 | 6 | *rd1* Untr. vs GW516 | *P* < 0.05 |
| | ZLN005 | 1.99 | ±0.56 | 6 | *rd1* Untr. vs ZLN005 | *P* < 0.0001 |

| B | | Mean % positive cells in ONL | Standard deviation (SD) | Sample (n) | Dunnett's multiple comparisons test | Significance(*p*) |
|---|---|---|---|---|---|---|
| Calpain-1 | *wt* Untr. | 0.27 | ±0.05 | 6 | *rd1* Untr. vs *wt* Untr. | *P* > 0.05 |
| | *rd1* Untr. | 0.40 | ±0.05 | 10 | | |
| | GW735 | 0.41 | ±0.04 | 6 | *rd1* Untr. vs GW735 | *P* > 0.05 |
| | Inolitazone | 1.00 | ±0.24 | 6 | *rd1* Untr. vs Inolitazone | *P* < 0.0001 |
| | GW516 | 0.70 | ±0.12 | 6 | *rd1* Untr. vs GW516 | *P* < 0.01 |
| | ZLN005 | 0.88 | ±0.23 | 6 | *rd1* Untr. vs ZLN005 | *P* < 0.0001 |

| C | | Mean % positive cells in ONL | Standard deviation (SD) | Sample (n) | Dunnett's multiple comparisons test | Significance(*p*) |
|---|---|---|---|---|---|---|
| Calpain-2 | *wt* Untr. | 0.51 | ±0.21 | 8 | *rd1* Untr. vs *wt* Untr. | *P* < 0.0001 |
| | *rd1* Untr. | 4.68 | ±0.50 | 9 | | |
| | GW735 | 4.08 | ±0.32 | 7 | *rd1* Untr. vs GW735 | *P* > 0.05 |
| | Inolitazone | 2.62 | ±0.56 | 5 | *rd1* Untr. vs Inolitazone | *P* < 0.0001 |
| | GW516 | 4.21 | ±0.42 | 4 | *rd1* Untr. vs GW516 | *P* > 0.05 |
| | ZLN005 | 2.94 | ±0.62 | 6 | *rd1* Untr. vs ZLN005 | *P* < 0.0001 |

**Supplemental Table 4. Effect of resveratrol or PJ34 on photoreceptor cell death (TUNEL assay) in the outer nuclear layer (ONL).**

| | | | Mean % positive cells in ONL | Standard deviation (SD) | Sample (n) | Dunnett's multiple comparisons test | Significance(*p*) |
|---|---|---|---|---|---|---|---|
| *rd1* | Untr. | | 3.55 | ±0.73 | 27 | *rd1* Untr. vs *wt* Untr. | *P* < 0.0001 |
| | Resveratrol | 10 µM | 3.74 | ±0.84 | 4 | *rd1* Untr. vs 10 µM Resveratrol | *P* > 0.05 |
| | | 20 µM | 3.35 | ±0.39 | 5 | *rd1* Untr. vs 20 µM Resveratrol | *P* > 0.05 |
| | | 100 µM | 3.87 | ±0.94 | 5 | *rd1* Untr. vs 100 µM Resveratrol | *P* > 0.05 |
| | PJ34 | | 2.83 | ±0.71 | 8 | *rd1* Untr. vs PJ34 | *P* < 0.01 |
| | Resveratrol + PJ34 | | 1.80 | ±0.19 | 5 | *rd1* Untr. vs Resveratrol + PJ34 | *P* < 0.0001 |
| C3H | Untr. | | 1.85 | ±0.59 | 14 | *wt* Untr. vs *rd1* Untr. | *P* < 0.0001 |
